# Cadmium Exposure Increases the Risk of Juvenile Obesity: A Human and Zebrafish Comparative Study

**DOI:** 10.1101/172346

**Authors:** Adrian J. Green, Cathrine Hoyo, Carolyn J. Mattingly, Yiwen Luo, Jung-Ying Tzeng, Susan Murphy, Antonio Planchart

**Affiliations:** Department of Biological Sciences and the, North Carolina State University, Raleigh, NC 27695; Center for Human Health and the Environment, North Carolina State University, Raleigh, NC 27695; Department of Statistics, North Carolina State University, Raleigh, NC 27695

## Abstract

**OBJECTIVE:** Human obesity is a complex metabolic disorder disproportionately affecting people of lower socioeconomic strata, and ethnic minorities, especially African Americans and Hispanics. Although genetic predisposition and a positive energy balance are implicated in obesity, these factors alone do not account for the excess prevalence of obesity in lower socioeconomic populations. Therefore, environmental factors, including exposure to pesticides, heavy metals, and other contaminants, are agents widely suspected to have obesogenic activity, and they also are spatially correlated with lower socioeconomic status. Our study investigates the causal relationship between exposure to the heavy metal, cadmium (Cd), and obesity in a cohort of children and a zebrafish model of adipogenesis.

**DESIGN:** An extensive collection of first trimester maternal blood samples obtained as part of the Newborn Epigenetics Study (NEST) were analyzed for the presence Cd, and these results were cross analyzed with the weight-gain trajectory of the children through age five years. Next, the role of Cd as a potential obesogen was analyzed in an *in vivo* zebrafish model.

**RESULTS:** Our analysis indicates that the presence of Cd in maternal blood during pregnancy is associated with increased risk of juvenile obesity in the offspring, independent of other variables, including lead (Pb) and smoking status. Our results are recapitulated in a zebrafish model, in which exposure to Cd at levels approximating those observed in the NEST study is associated with increased adiposity.

**CONCLUSION:** Our findings identify Cd as potential human obesogen. Moreover, these observations are recapitulated in a zebrafish model, suggesting that the underlying mechanisms may be evolutionarily conserved, and that zebrafish may be a valuable model for uncovering pathways leading to Cd-mediated obesity in human populations.

## INTRODUCTION

The prevalence of obesity has more than doubled among children and more than tripled among adolescents in the last 30 years ^1,2^. While obesity prevalence has plateaued overall in the last two years, the disparities in the prevalence of obesity in children of lower socioeconomic status (SES) and racial/ethnic minorities appear to be widening ^3-5^. Genetic predisposition and energy imbalance, where caloric input exceeds energy output, are implicated in obesity; however, these factors alone cannot explain the disproportionate incidence of obesity in lower SES populations. The increased use of organic and inorganic chemicals for a wide range of applications in the last century has been paralleled by increases in the body burden of environmental pollutants, many of them endocrine disruptors. In animal models, *in vitro* and in humans, many of these chemicals have been associated with lipid accumulation and progressive cardiometabolic dysfunction. However, these data have been difficult to interpret and use to recommend public action, as the specificity of the associations between many of these chemicals and the cardiometabolic disease risk phenotype has not been demonstrated, and the doses of exposure in model systems are often at or above human occupational levels.

Cadmium (Cd) is a ubiquitous environmental contaminant ranked seventh on the list of toxicants of concern by the Agency for Toxic Substances and Disease Registry (ATSDR)^6^. Two to three decades leading up to the 1970s saw a rapid increase in the use of Cd in the manufacture of fertilizer and nickel-cadmium batteries, that paralleled an increase in blood Cd concentrations in the US population ^7-10^. Major sources of human exposure include ingestion of foods contaminated with Cd, cigarette smoke, and breathing contaminated air in occupational settings or in neighborhoods near contaminated industrial facilities. The mechanisms by which Cd elicits toxicity are not entirely clear, although induction of oxidative stress has been implicated. Understanding the connection between exposure and Cd-mediated outcomes may be further complicated by its long half-life, estimated to be between 10 and 45 years, in the kidney, liver, lung and pancreas^11,12^. Cd is a known human carcinogen and is associated with respiratory, renal, neurological, and bone disorders. In addition, some studies^13-15^, including reviews^12,16-18^, but not others^19,20^ link lower levels of Cd to cardiovascular and metabolic diseases; however, these associations are limited to adults.

Epidemiological and animal studies over the past 15 years have demonstrated that *in utero* and neonatal environmental exposures alter programming of endocrine systems involved in growth, energy metabolism, adipogenesis, appetite, and glucose-insulin homeostasis of the developing fetus^21-25^. Cd exposure has been associated with lower birth weight^26-28^, a phenomenon known to be a persistent risk factor for accelerated adiposity gain in young children, which has been linked to cardio-metabolic impairment in adulthood^29-35^. Exposures occurring during critical developmental windows have been shown to stably alter the function of target organ systems, and initiate processes that increase the risk of cardiometabolic diseases later in life^29,36^. Currently cohort data linking low-level prenatal Cd exposure to cardiometabolic outcomes are limited and derive from studies with short follow-up^37-39^. Thus, it remains unclear whether early indications of metabolic dysfunction that have been associated with developmental exposure to Cd persist into middle childhood or adulthood. Furthermore, because prenatal Cd exposure also disproportionately affects lower SES strata, disentangling the contributions of Cd from competing risk factors including physical activity, dietary patterns, and other non-chemical stressors, has thus far not been possible^40^. Additional models are needed to isolate the effects of early developmental exposure to Cd on metabolic indicators.

Zebrafish (*Danio rerio*) is a powerful model system for toxicological research^41,42^. Its genome is sequenced and its conservation with humans is facilitating mechanism-based understanding of chemical effects on diverse human conditions^43^. Its experimental strengths include its small size, high fecundity, availability of transgenic lines for live imaging of complex physiological processes, embryonic transparency, experimental tractability, and conserved but simplified anatomy^41,42^. Zebrafish larvae and adults are semitransparent and offer unique opportunities to study the effects of environmental exposures on adipogenesis and metabolic function *in vivo*^44^., Adipose tissue is recognized as a dynamic endocrine organ that plays a critical role in regulating metabolic homeostasis^45^, in addition to storing excess fat. Adipose tissue is first detected in zebrafish at about two weeks post-fertilization, embryonic and early larval stages are sensitive to compounds that modulate fat metabolism^44,46-48^. The deposition and mobilization of lipid within zebrafish adipose tissue can be altered by nutritional manipulation, suggesting that` energy storage functions of adipose tissue are conserved between zebrafish and mammals^49^. In addition, gene expression studies on unfractionated zebrafish adipose tissue show shared pathophysiologic pathways indicating that zebrafish studies involving adipogenesis and metabolic function may be directly translatable to humans^49,50^.

Here, we present human data linking prenatal Cd exposure to obesity in children at age five years, and demonstrate that this effect is recapitulated in juvenile zebrafish exposed to Cd during the larval stage. Despite the likely presence of confounders in the human data, our findings in zebrafish, in which the exposure profile is strictly controlled, demonstrate for the first time that Cd may be a human obesogen, and that prenatal human exposure to Cd likely initiates a cascade of molecular events leading to increased adiposity.

## MATERIALS AND METHODS

### Study participants

Study participants were pregnant women enrolled in the Newborn Epigenetic STudy (NEST), a prospective cohort study of women and their offspring enrolled from 2009 to 2011 from six prenatal clinics in Durham County, North Carolina. Participant accrual procedures were previously described^51,52^. Briefly, inclusion criteria were: age 18 years or older, pregnant, and intention to use one of two participating obstetric facilities in Durham County for delivery. Exclusions were: plans to relinquish custody of the index child, move states in the subsequent three years, or an established HIV infection. In the 18-months beginning April, 2009, 2,548 women were approached and 1,700 consented (66.7% response rate). The present analyses are limited to the first 319 infant-mother pairs in whom we measured first trimester blood Cd, arsenic (As) and lead (Pb). Maternal race, smoking status, BMI before pregnancy, parity, delivery route, and education were comparable in the 319 infant-mother pairs included in this study and the remainder of the cohort (p>0.05). The study protocol was approved by the Duke University Institutional Review Board.

### Data and specimen collection

Participants completed a self-or interviewer-administered questionnaire at the time of enrollment that included social and demographic characteristics, reproductive history, lifestyle factors, and anthropometric measurements. At study enrollment, maternal peripheral blood samples were collected; the mean gestational age at maternal blood draw was 12 weeks. Blood aliquots were prepared and stored at −80°C.

### Measurement of cadmium

Prenatal Cd blood levels were measured in whole blood as nanograms per gram (ng/g; 1000ng/g=1035ng/μl) using well-established solution-based ICP-MS methods^53-56^. Procedures were described previously^26^. Briefly, frozen maternal blood samples were equilibrated at room temperature, homogenized with a laboratory slow shaker (GlobalSpec, East Greenbrush, NY) and ∼0.2 mL aliquots were pipetted into a trace-metal-clean test tube and verified gravimetrically to ±0.001mg using a calibrated mass balance. Samples were spiked with internal standards consisting of known quantities (10 and 1 ng/g, respectively) of indium (In) and bismuth (Bi) (SCP Science, USA), used to correct for instrument drift. The solutions were then diluted using water purified to 18.2 MΩ/cm resistance, hereinafter referred to as Milli-Q water (Millipore, Bedford, Mass., USA) and acidified using ultra-pure 12.4 mol/L hydrochloric acid to result in a final concentration of 2% hydrochloric acid (by volume). All standards, including aliquots of the certified NIST 955c, and procedural blanks were prepared by the same process.

Cd concentrations were measured using a Perkin Elmer DRC II (Dynamic Reaction Cell) axial field ICP-MS at the University of Massachusetts-Boston^53-56^. To clean sample lines and reduce memory effects, sample lines were sequentially washed using Milli-Q water for 90 seconds and a 2% nitric acid solution for 120 seconds between analyses. Procedural blanks were analyzed within each block of 10 samples, to monitor and correct for instrument and procedural backgrounds. Calibration standards used to determine metal in blood included aliquots of Milli-Q water, and NIST 955c SRM spiked with known quantities of each metal in a linear range from 0.025 to 10 ng/g. Standards were prepared from 1000 mg/L single element standards (SCP Science, USA). Method detection limits (MDLs) were calculated according to the two-step approach using the t_99_S_LLMV_ method (USEPA, 1993) at 99% CI (t=3.71). The MDLs yielded values of 0.006, 0.005, and 0.071 μg/dL, for Cd, Pb, and As, respectively. Limits of detection (LOD) were 0.002, 0.002, and 0.022 μg/dL, for Cd, Pb and As, respectively, and limits of quantification (LOQ) (according to Long and Winefordner, 1983) were 0.0007, 0.0006, and 0.0073 μg/dL for Cd, Pb, and As, respectively. The number of samples below the LOD for Cd, Pb, and As were 2, 2, and 1, respectively.

### Statistical analyses

Childhood obesity at age five was defined by the weight-for-height z score (WHZ)^57^. Children with WHZ scores greater than 85% of their same sex peers at age five were classified as overweight/obese. Logistic regression was implemented to evaluate the association between childhood obesity and the concentration of Cd, adjusting for other co-occurring metals (Pb and As) in maternal blood, maternal smoking (never, quit during pregnancy, pregnant smoker), breastfeeding (over three months or less), and sex of child. To reduce bias related to episodic growth acceleration, we additionally adjusted for child weight trajectory from birth to 36 months. These growth trajectories were computed as growth curves for each child, and functional principal component analysis (FPCA) was implemented to summarize growth curves. In the final model the top two FPCs, which explain 95% of the variability in the original growth curves were included as covariates in the regression model. Similar to PCA (which aims to extract orthogonal PCs that retain maximal amount of variation in the original variables by estimating the eigenvalues and eigenvectors of the sample variance-covariance matrix), FPCA aims to obtain orthogonal functional PCs that retain the maximal amount of variation in the original weight curves by estimating the eigenvalues and eigenfunctions of the sample variance-covariance function.

### Zebrafish husbandry and embryo collection

Wildtype (AB) zebrafish were maintained in a zebrafish facility at NC State University according to standard protocols,^58^ and in conformity with guidelines of the NC State Animal Care and Use Committee (ACUC), which also approved all animal experiments reported.. Briefly, adults were maintained at 28.5° C and a 14/10-hour light/dark cycle, and fed a standard diet twice daily. Spawning took place at a ratio of three females to one male; embryos were collected every 30 minutes and scored for viability prior to use in downstream applications.

### Radioassay to assess cadmium uptake by larval zebrafish

To assess total body concentrations of Cd in zebrafish, triplicate groups of zebrafish embryos (n=25/group) were exposed from four hours post-fertilization (hpf) to seven days post-fertilization (dpf) to 60 μg/L of Cd in the form of CdCl_2_ in 0.5x embryo media (E2), spiked with ^109^Cd as a tracer (1592 Bq μg^−1^). Solutions were replaced daily during the course of the experiment. Larval uptake of Cd was monitored daily beginning at three dpf by measuring radioactive decay corrected for background activity. Briefly, larvae were washed three times with five ml of Cd-free, non-radioactive 0.5x E2 media followed by transfer to clean scintillation vials in two mL of the final wash. An additional two mL of the final wash were transferred to a second clean scintillation vial to measure background activity. The radioactivity uptake was measured using a Wallac Wizard 1480 Gamma counter.

### Cadmium exposure

Stock solutions of CdCl_2_ ([Cd], 99.99% purity; Sigma-Aldrich, MO) were made at 60 parts per million (1000x), in Milli-Q water. Zebrafish embryos were collected as described and exposed to 60 parts per billion (ppb) Cd in 0.5X embryo media^58^ from four hpf to seven dpf at a density of 10 embryos/mL with daily replacement, and fed beginning at five dpf. After removal of Cd, larvae were raised for lipid content analysis at one and two months post-fertilization.

### Lipid analysis

The vital dye, Nile red, was used to stain lipids in juvenile zebrafish (one and two months post-fertilization), which allows repeated analysis of the same individual to assess amount and location of lipid droplets over time^49^. A 1.25 mg/mL stock solution was made in Milli-Q water. Immediately before use, a working solution was made by diluting 10 μL of the stock solution into 25 mL of aquarium system water to provide a final concentration of 0.5 μg/mL. Live zebrafish were stained in the dark for 30 minutes at 28°C^44,49^. Fish were removed from the Nile red solution and anesthetized in aquarium system water containing 0.25 mg/mL phosphate buffered (pH 7) Tricaine-S (Western Chemical, Ferndale, WA).

### Imaging and quantitative analysis

Nile red-stained zebrafish were imaged using a Leica MZ FLIII fluorescence stereomicroscope. Images were analyzed using Fiji^59^. Color thresholding was used to select Nile red-containing sections by setting the hue value at 20-50. Background fluorescence was removed by setting a minimum brightness threshold of 120. Remaining fluorescence was selected and analyzed using the measure tool^44,60,61^. To account for differences in body size, fluorescence was normalized by taking the ratio of fluorescence to the dorsal-ventral height at the point where the anal fin attaches anteriorly to the body ^62^.

## RESULTS

### Study subjects

The distributions of first trimester blood Cd concentrations were compared by social and demographic characteristics of the mother-child pairs (Table 1). African Americans comprised 35% of the study population while Whites, Hispanics and Others comprised 30%, 32% and 4%, respectively. Nearly two thirds were younger than 30 years; approximately half had at least a high school education level, and reported a household income of at least $25,000 per year. Seventy-three percent were married or living with a partner. Fifteen percent of mothers reported smoking during pregnancy and 55% were overweight, obese, or extremely obese (29%, 15%, or 11% respectively). The majority of offspring (89%) had a birth weight within normal range (2.5 to 4 kg) and 88% were born at term. Blood Cd and Pb concentrations did not vary by maternal age, obesity, gestational age at delivery, or by sex and birth weight of offspring. However, blood levels of these heavy metals were higher among infants born to African Americans, Asians and Hispanics compared to Whites (p=0.03), smokers (p=0.01), and those who were obese before pregnancy (p=0.02). These factors were considered as potential confounders.

**Table 1.** Description of characteristics for study participants.

| Category |  | N | Cadmium (ng/g)<br>quantile: median<br>[IQR*] | Lead (ng/g)<br>quantile: median<br>[IQR*] |
| --- | --- | --- | --- | --- |
| Maternal age<br>(in years) | <30 | 182 | 0.1 [0, 0.2] | 1.6 [0, 3.5] |
|  | 30<35 | 76 | 0.1 [0, 0.2] | 1.5 [0.5, 2.8] |
|  | 35+ | 56 | 0.1 [0, 0.1] | 2.0 [0.4, 5] |
| Maternal<br>educational levels | Less than high school<br>or high school | 162 | 0.1 [0, 0.3] | 2.1 [0, 4.1] |
|  | College | 151 | 0.1 [0, 0.1] | 1.4 [0.4, 2.9] |
|  | Graduate degree | 1 | 0.2 [0.2, 0.2] | 1.7 [1.7, 1.7] |
| Ethnic composition | White | 96 | 0.1 [0, 0.1] | 1.3 [0.4, 2.4] |
|  | Black | 108 | 0.1 [0, 0.3] | 1.6 [0, 3.3] |
|  | Hispanic | 98 | 0.1 [0, 0.2] | 2.2 [0, 4.9] |
|  | Other | 12 | 0.1 [0, 0.2] | 2.7 [0.7, 5.3] |
| Cigarette smoking | Never Smoked | 228 | 0.1 [0, 0.2] | 1.5 [0.4, 3.5] |
|  | Smoking during<br>pregnancy | 46 | 0.3 [0, 0.4] | 1.7 [0, 3.2] |
|  | Smoking prior to<br>pregnancy only | 40 | 0.1 [0, 0.2] | 1.7 [0, 2.6] |

\*\*IQR: interquartile range

### Associations between first trimester cadmium and obesity

Maternal first trimester blood Cd concentrations were 0.3 ng/g of blood weight (IQR0.1-0.7), i.e. 0.03μg/dL, which is comparable to the US population^63^. Higher prenatal Cd levels were associated with higher obesity risk at five years of age (Table 2). The effect of Cd (β=3.211, se=1.33, p=0.03) corresponds to a ∼25-fold increase in obesity odds at age five for every one ng/g increase in blood weight of Cd. These analyses were adjusted for sex, cigarette smoking, exposure to Pb and As, and the first two functional principal components of growth trajectories. Figure 1 also shows the increase in the magnitude of the adjusted associations between first trimester Cd exposure and obesity at each month with increasing age, until 30 months when it plateaus, indicating that Cd-associated obesity is likely sustained, at least in childhood. Additional adjustment for pre-pregnancy obesity did not alter these associations.

**Table 2.** Adjusted regression coefficients for associations between cadmium exposure and obesity parameters, in children at age 4-5 years*.

| Parameter | Regression Coefficient | Std. Error | p-value |
| --- | --- | --- | --- |
| Intercept | 3.97 | 2.78 | 0.395 |
| Functional principal components for growth trajectories** | 1.22 | 0.42 | 0.004 |
| Functional principal components for growth trajectories | 1.40 | 1.50 | 0.353 |
| Prenatal blood Cd concentrations | 2.91 | 1.34 | 0.030 |
| Prenatal As concentrations | -13.80 | 7.47 | 0.065 |

\*Model adjusted for Pb concentrations, sex, breastfeeding for at least 3 months.
\*\*Functional principal components summarize growth trajectories from birth to age 3 years and are mutually exclusive.

**Figure 1.**
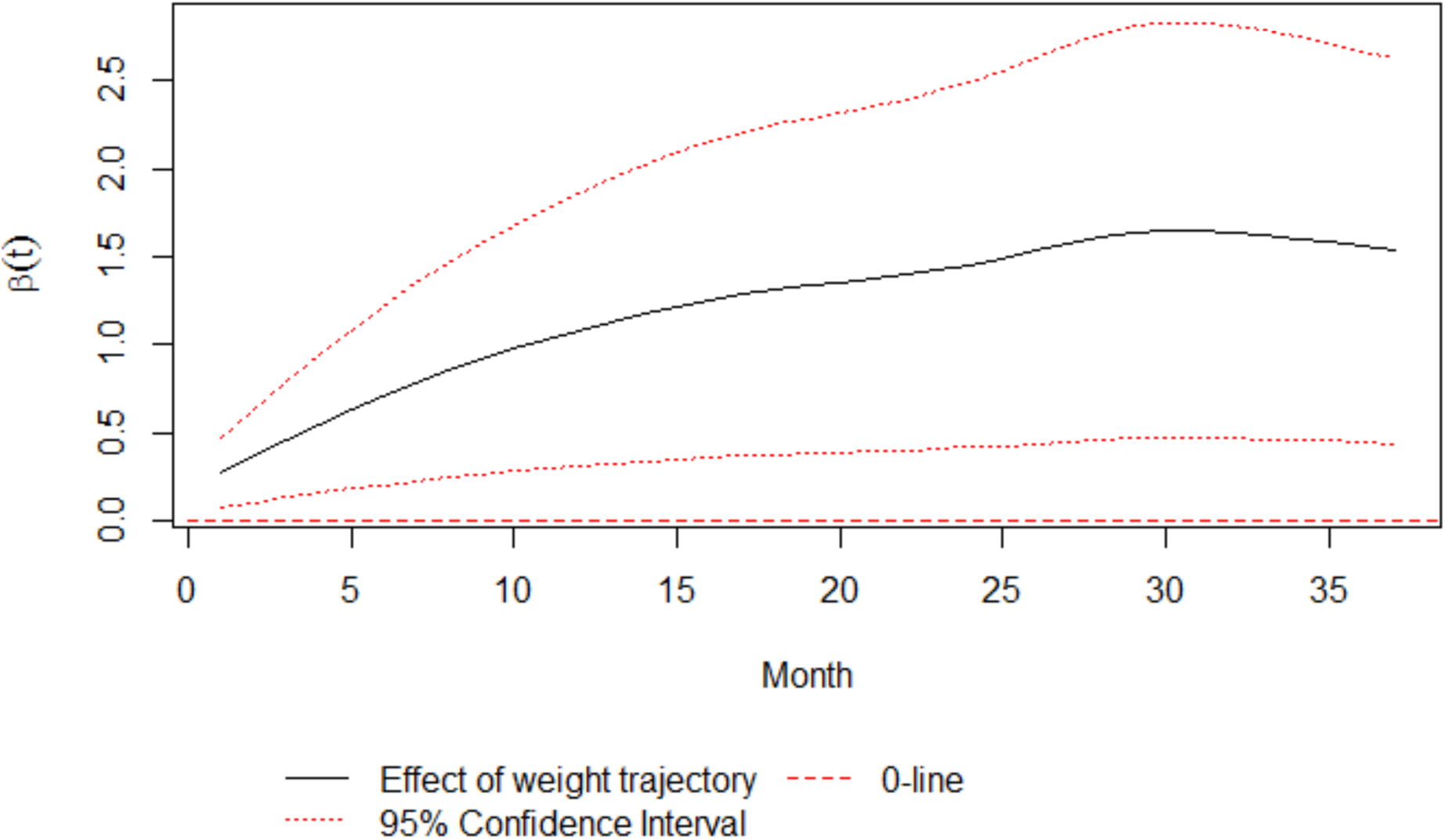
Effect of weight trajectory (via the first FPC) on obesity risk at age five. The solid line indicates the effect of child weight by month via the first FPC on obesity risk at age five; the flanking dashed lines represent the 95% simultaneous confidence band of the weight effect, accounting for multiple comparisons of all months; the dotted line indicates zero effects. The simultaneous confidence band lies above zero, indicating a significant, positive effect of child weight on obesity risk at age five. The solid line also suggests that the magnitude of the weight effect increases over time.

### Cadmium uptake by larval zebrafish

Larval zebrafish began to accumulate measurable amounts of Cd from three dpf onward (Figure 2). The delay in Cd uptake correlated with the presence of the chorion, an embryonic membrane surrounding the developing embryo that typically ruptures at or about 48 hpf. Beginning at three dpf, Cd accumulation was approximately linear, and at seven dpf the total body burden of Cd reached 0.54 ng ± 0.1 ng/larvae. On average, a seven dpf larval zebrafish weighs 1.4 mg (Hu et al., 2000); by extrapolation, this equates to 386 ng Cd per gram of larvae. Since Cd burden is commonly reported as a serum concentration, we used the Cd toxicokinetic model proposed by Kjellström and Nordberg^64^ to estimate a larval serum concentration. This model estimates that 0.06% of the total body burden of Cd can be found in the serum; therefore, the calculated serum concentration per larvae is 0.23 ng/g, in agreement with the values observed in the NEST cohort.

**Figure 2.**
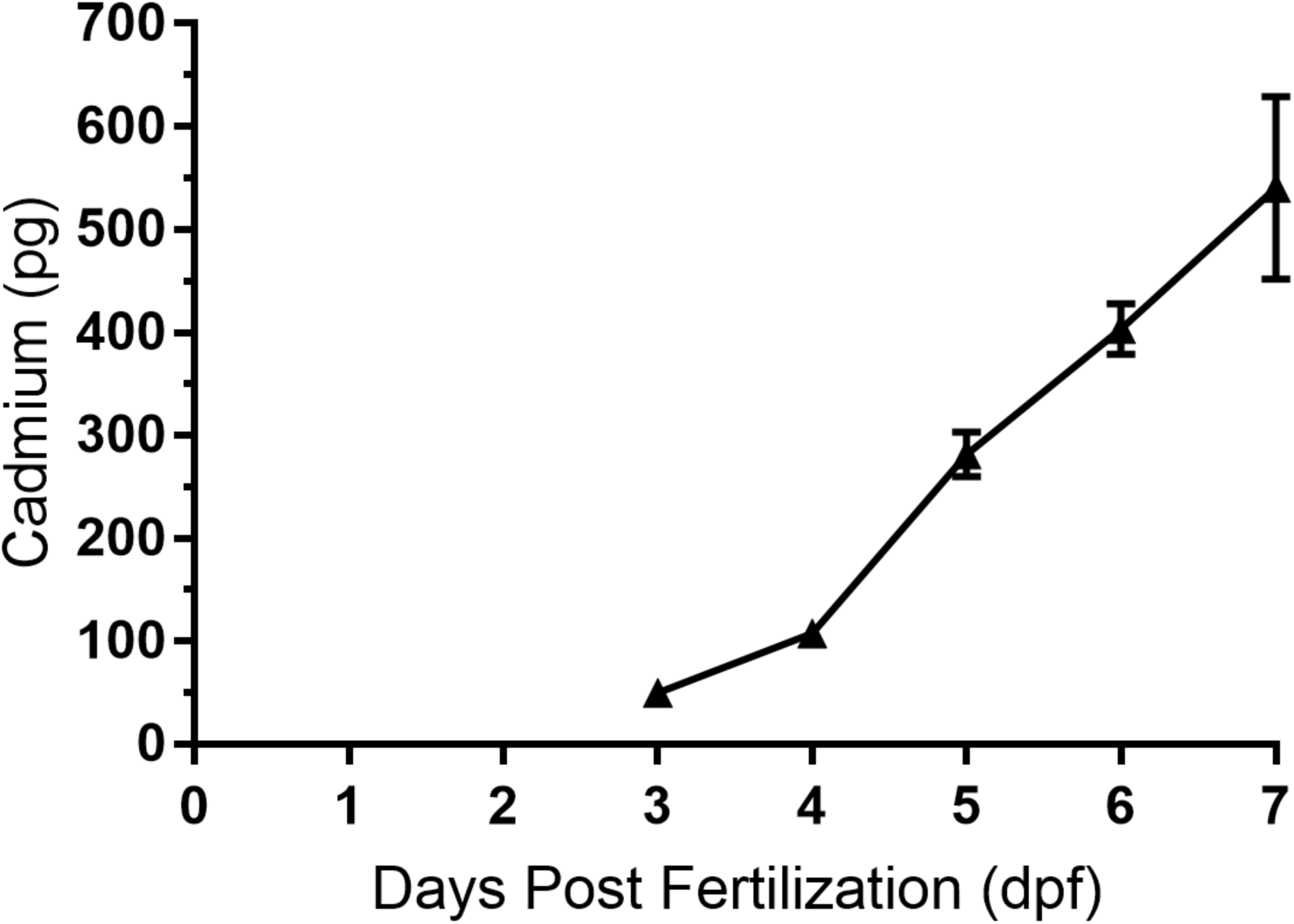
Total cadmium uptake during zebrafish development. Total internal Cd was measured as described after zebrafish embryos were exposed continuously from four hpf to seven dpf to Cd spiked with ^109^Cd. Measurements began at three dpf after hatching from the chorion, which provides a significant barrier to Cd uptake. Measurements are mean±SEM.

### Cadmium-induced juvenile lipid accumulation

Zebrafish undergo rapid development, with free-feeding larvae emerging after five dpf. However, a prolonged juvenile period of approximately three months follows, resulting in sexually mature adults at about 3-3.5 months post-fertilization. Zebrafish exposed to 60 ppb Cd during embryonic/larval development had significantly increased lipid accumulation at one and two months post-fertilization as seen in size-adjusted Nile red fluorescence following exposure from four hpf to one week post-fertilization (Figure 3, p < 0.05). This increase in Nile red fluorescence was not seen at 3.5 months post-fertilization (data not shown) at which point the Nile red fluorescence was significantly decreased in the Cd-exposed group vs controls (p < 0.01). These data indicate that limited (developmental) exposure to Cd results in increased lipid accumulation in juvenile zebrafish, which persists throughout the pre-and peri-pubertal stages but likely reverses at or before the onset of sexual maturity in the absence of continuous exposure.

**Figure 3.**
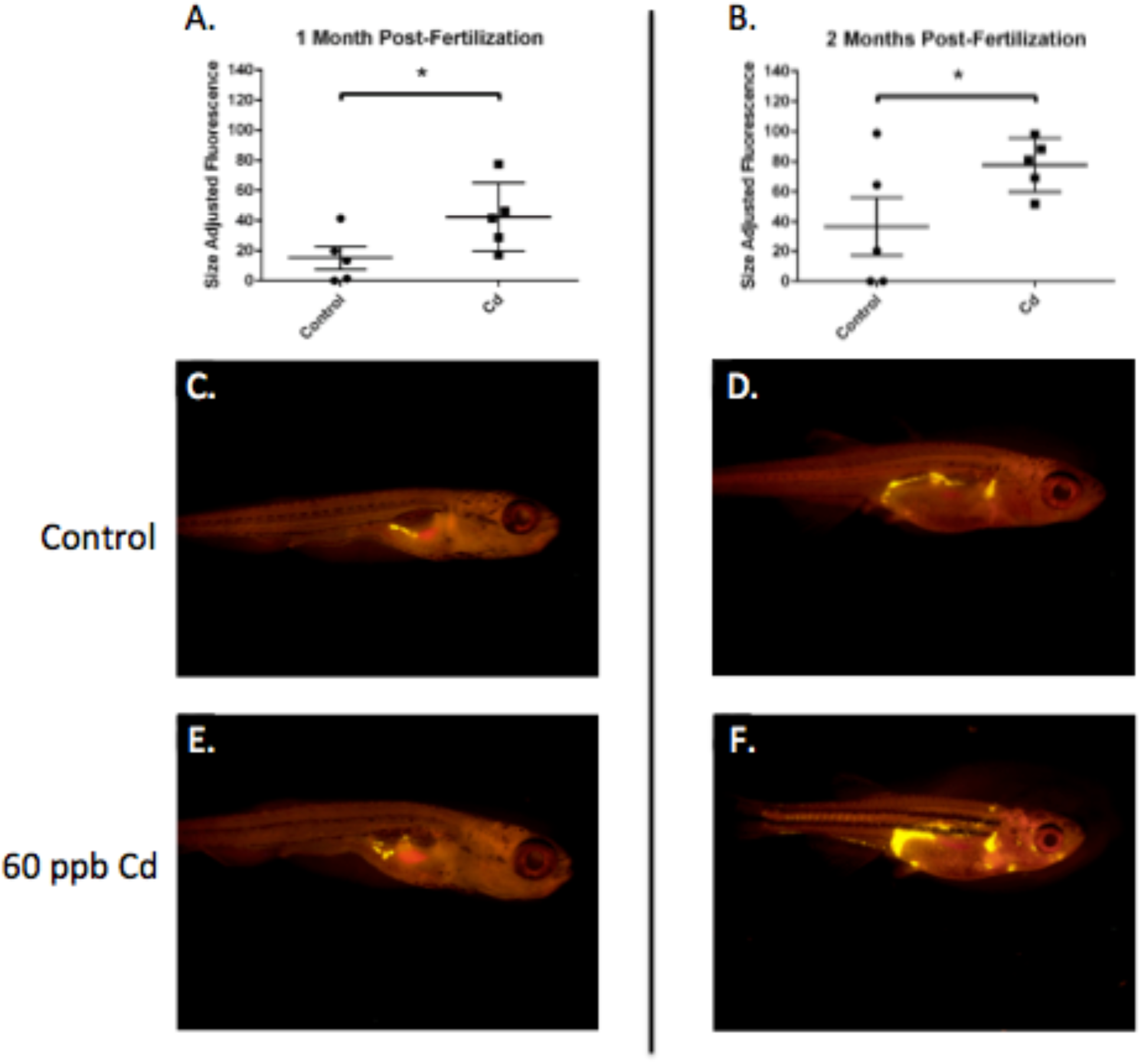
Developmental exposure to cadmium increases lipid deposition in juvenile zebrafish. Nile red fluorescence was significantly greater in zebrafish larvae exposed to 60 ppb vs. water controls at one (A) and two (B) months post-fertilization (p<0.05). Representative live images of Nile red staining are shown for control (C, D) and Cd-exposed (E, F) zebrafish at one-and two-months post-fertilization, respectively.

## DISCUSSION

Although genetic predisposition and energy imbalance, where energy input exceeds output, are established risk factors fueling the obesity epidemic in children, caloric excess and physical inactivity alone fail to fully account for the magnitude and the steep trajectory followed by the obesity epidemic^65^. A growing consensus suggests that exposure to some lipophilic or metalloid contaminants is obesogenic; the most studied are persistent organic compounds such as polychlorinated bisphenyls^66^, and metalloids such as arsenic^67-70^. However, the obesogenic potential of ubiquitous inorganic metals, including Cd, is unclear.

We evaluated associations between prenatal Cd exposure and obesity in children, and determined the plausibility of this relationship in a controlled experimental zebrafish model. After adjusting for cigarette smoking, sex, breastfeeding and co-occurring metals (Pb and/or As), we found persistent associations between prenatal Cd exposure and increased risk of obesity from birth to age five years. Our data also suggest that these children were also more likely to have steeper growth trajectories between birth to age five years. In support of this association, we also found that zebrafish exposed developmentally to Cd exhibited similar concentrations as those found in humans at similar developmental stages. Furthermore, these fish went on to exhibit significantly higher lipid accumulation as juveniles, when compared to unexposed controls. Surprisingly, lipid accumulation plateaued at or near the onset of sexual maturity. Although similar data observations are suggested in human data, follow-up is short and sample sizes small as evidenced by the wide confidence intervals. However, if similar plateauing of obesity risk were replicated in larger studies, these findings would support the intriguing possibility that, without postnatal exposure, Cd-associated obesity may in fact be transient.

To our knowledge, our study represents the first direct measure of association between prenatal Cd exposure and increased obesity risk in children, the results of which are supported by similar findings in an evolutionarily related model organism. Whether Cd is measured in biological materials that reflect long term chronic exposure, such as toe nails or urine or in blood, reflecting shorter term, concurrent exposure, data linking elevated Cd levels to obesity related cardiometabolic diseases among adults are inconsistent^13-15,12,16-18,19,20^. However, in early life, exposure to Cd is consistently associated with lower birth weight^26,27,71-73^, although the few studies that have examined the association between prenatal Cd and growth^73^ found that maternal Cd was associated with lower head circumference, height and weight. Reasons for inconsistent findings are unclear although differences in exposure dose, i.e., circulating concentration, could be a factor, which may depend on the source of exposure. Cd doses that are ingested or inhaled from contaminated air or dust are likely higher than levels in contaminated grains, which form only a fraction of the total diet. Inconsistent findings could also be due to co-exposure to other metals, which together with Cd, may have antagonistic effects, e.g., selenium. Differences could also be due to inadequate control for confounding by socioeconomic status, which in turn may influence not only dietary factors but also residing in geographic locations of higher exposure^74^. In zebrafish exposed only to Cd, limited to the human-equivalent periconceptional and early prenatal period and the elimination of socioeconomic effects, Cd exposure was associated with lipid accumulation. Whether the plateauing effect is sustained into puberty and beyond is still unknown.

Mechanisms linking low dose Cd exposure and subclinical cardiometabolic dysfunction are unclear; however, single metal analysis in adults suggests that blood Cd below reportable levels of 0.5 μg/dL was associated with elevated glucose^75-79^, higher blood pressure, presumably via kidney dysfunction ^80,81^, and oxidative stress^82^, which depletes antioxidants^83,84^. In autopsy specimens, higher liver Cd levels were associated with hypertension^85^. In mice and *in vitro*, early Cd exposure increased inflammation, oxidative stress, and blood pressure, doubled adipocyte numbers^86^, and lowered the expression of lipid synthesis genes^87^; thus obesity could result *directly* from this increased capacity for lipid storage. In these model systems, early Cd exposure also dysregulated the release of chemokines, leptin and adiponectin^86,87^ leading to insulin resistance later in life^88^. As these chemokines are involved in appetite regulation and energy expenditure^89-91^, cardiometabolic dysfunction indicators may also result *indirectly* via altered satiety responsiveness and increased caloric intake. Disentangling these possibilities will be critical in the future, to guide intervention efforts aimed at reducing Cd-related cardiometabolic dysfunction.

A major strength of our study is the ability to demonstrate in humans and in zebrafish that Cd increases lipid accumulation, leading to obesity, and associations are free from the influence of co-exposure to other metals and socioeconomic factors. However, our study had a limited sample size as evidenced by the wide confidence bands. While the sample size was adequate to demonstrate significant associations in overall analyses, we were under-powered to examine sex differences in children; Cd exposure effects may vary by sex. In addition, although prospective, children were followed from birth to age five years, and without serial specimens, the effects of postnatal exposure could not be disentangled in children. However, zebrafish that were exposed only “prenatally” had significantly higher lipid accumulation that the unexposed controls, suggesting that postnatal exposure did not unduly influence our findings in children. Moreover, the extent to which Cd-related obesity will be maintained after age five years is unknown. Zebrafish that were followed until sexual maturity exhibited reduced lipid accumulation.

Despite these limitations, our data support the causal association between *in utero* exposure to Cd and obesity at age five years. Larger studies are required to confirm these findings and determine Cd effects vary by sex.

## ACKNOWLEDGEMENTS

We thank participants of the NEST project. We also acknowledge Stacy Murray, Kennetra Irby, Siobhan Greene and Anna Tsent for recruiting NEST participants, Carole Grenier and Erin Erginer for technical assistance, Dr. David Buchwalter for assistance with the Cd uptake assay, Carson Lunsford for assistance with the zebrafish lipid assay, and David C. Cole for zebrafish husbandry. AJG was supported by the Ruth L. Kirschstein National Research Service Award Institutional Training grant number T32ES007046. This publication was supported in part by NIH under award numbers P30ES025128 (CH, CJM, AP) and R01ES016772 (CH), and by a generous donation from Howard and Julia Clark.

